## Supplementary figures and images for "Genome-wide functional analysis reveals key roles for kinesins in the mammalian and mosquito stages of the malaria parasite life cycle"

### S1

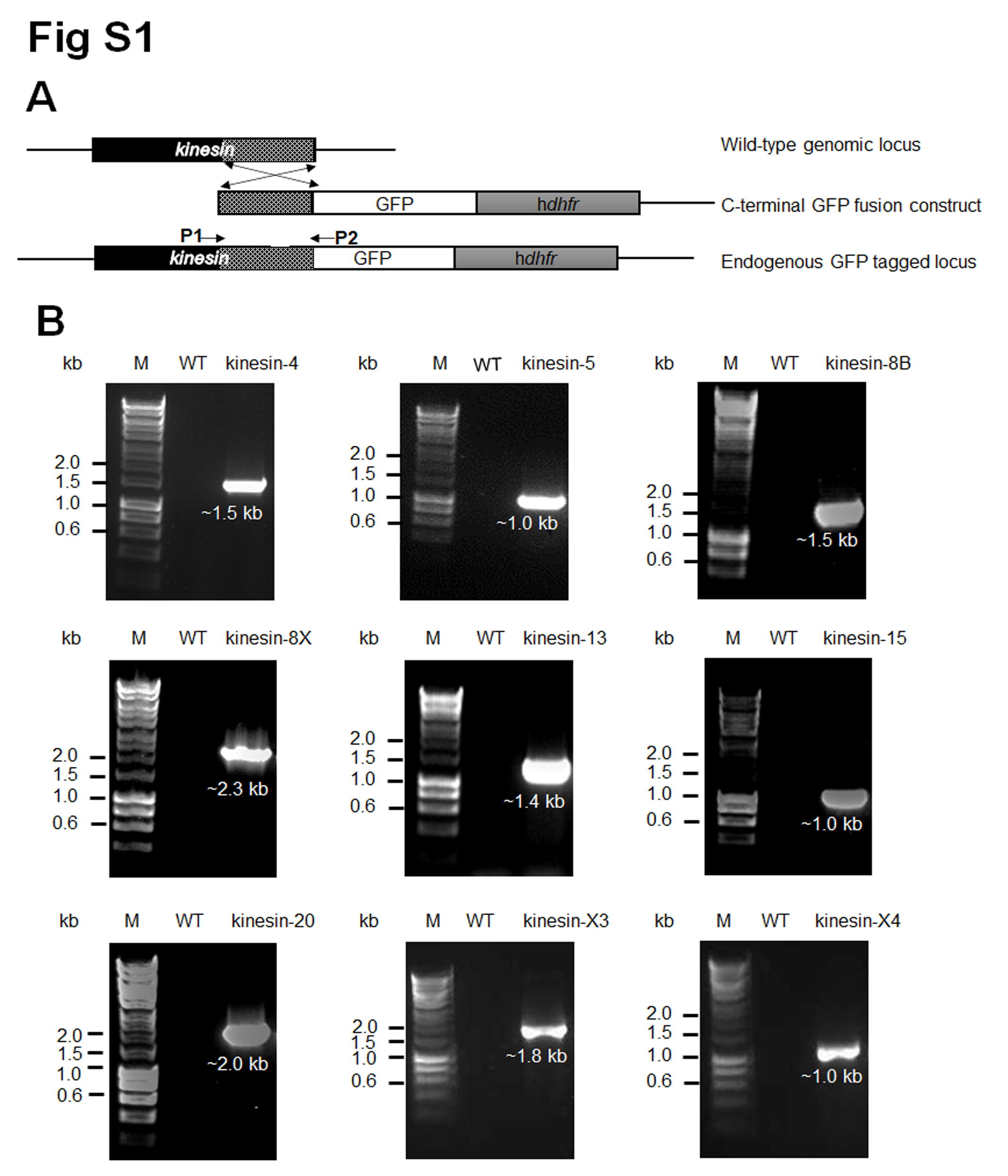

### S2

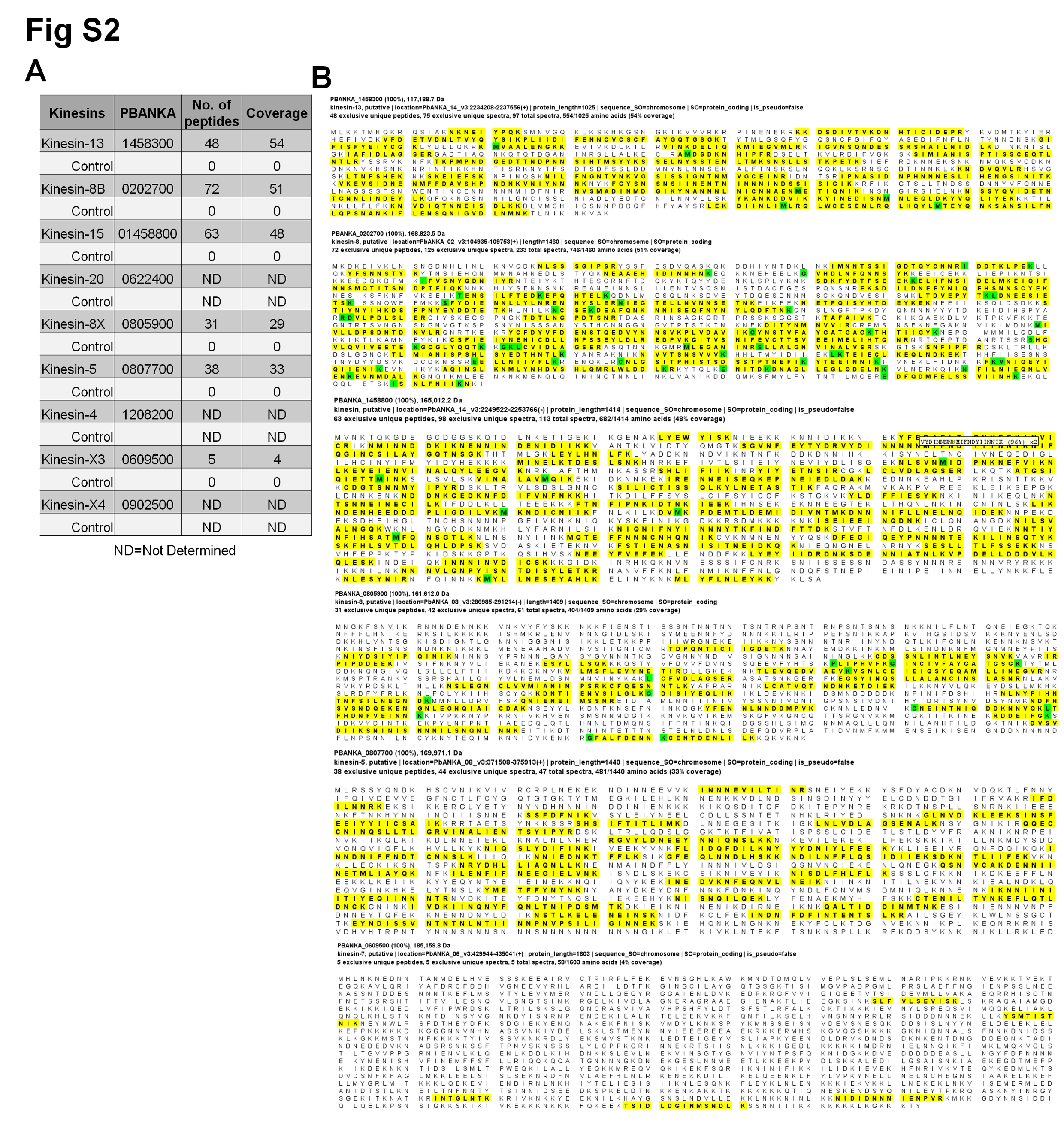

### S3

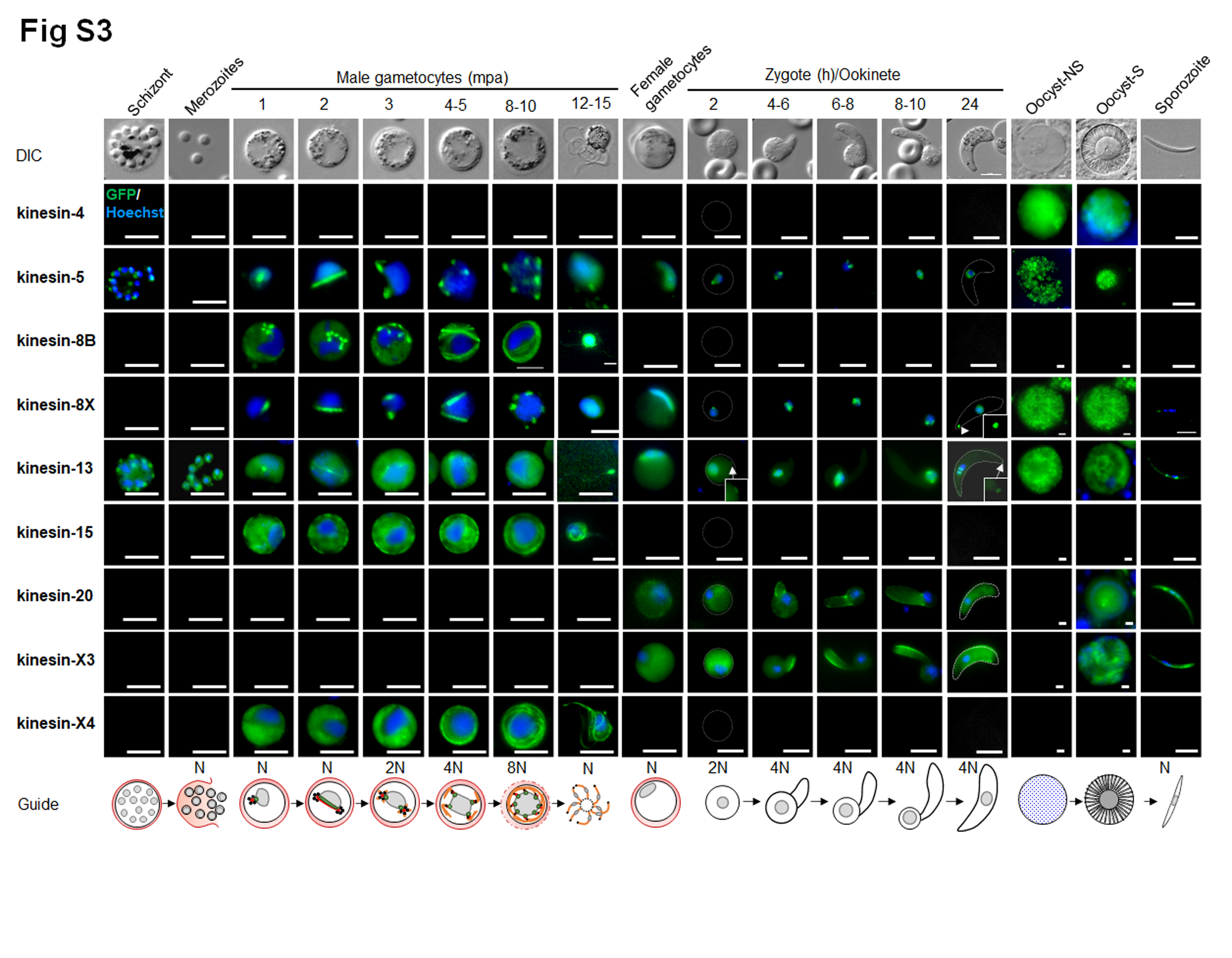

### S4

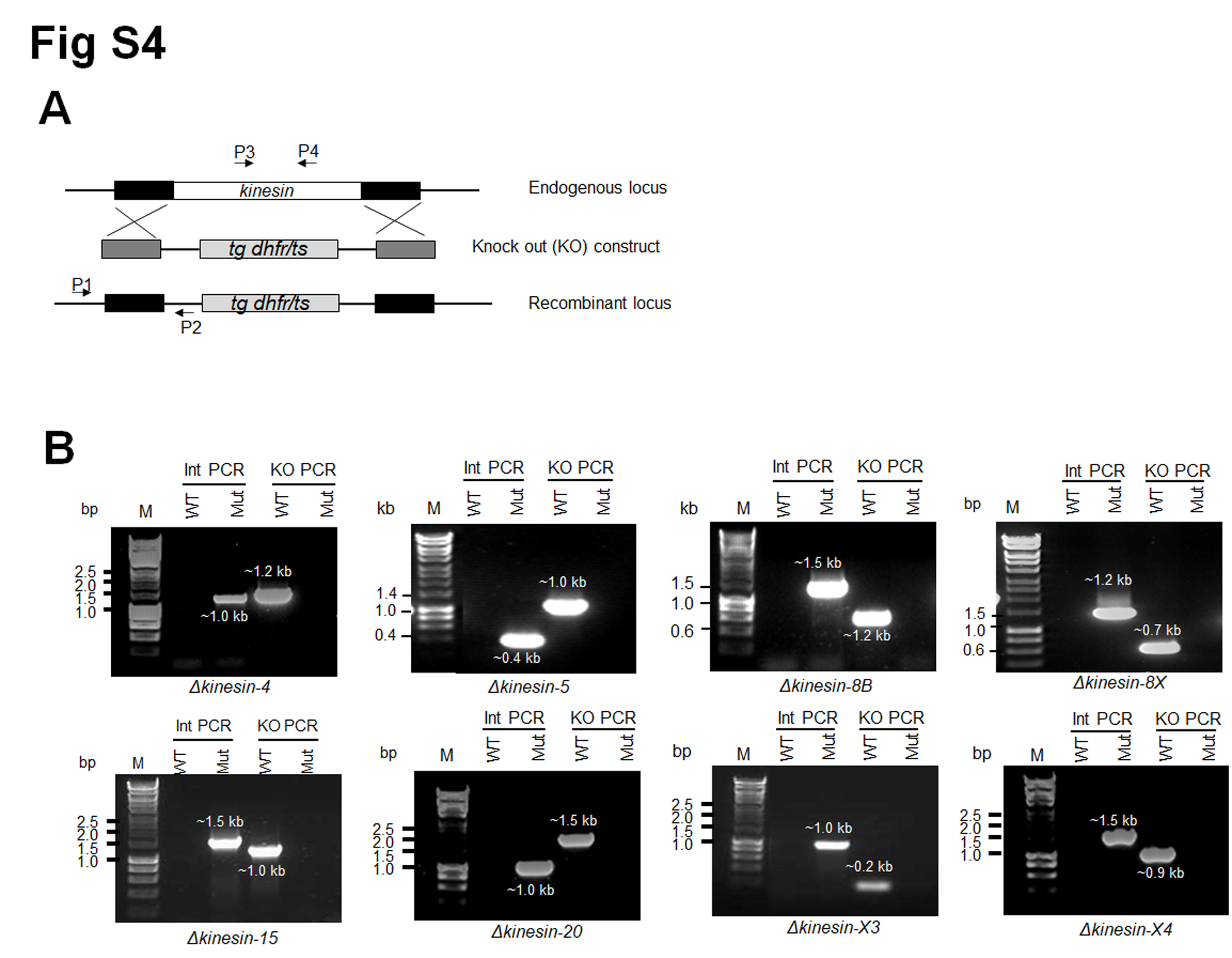

### S5

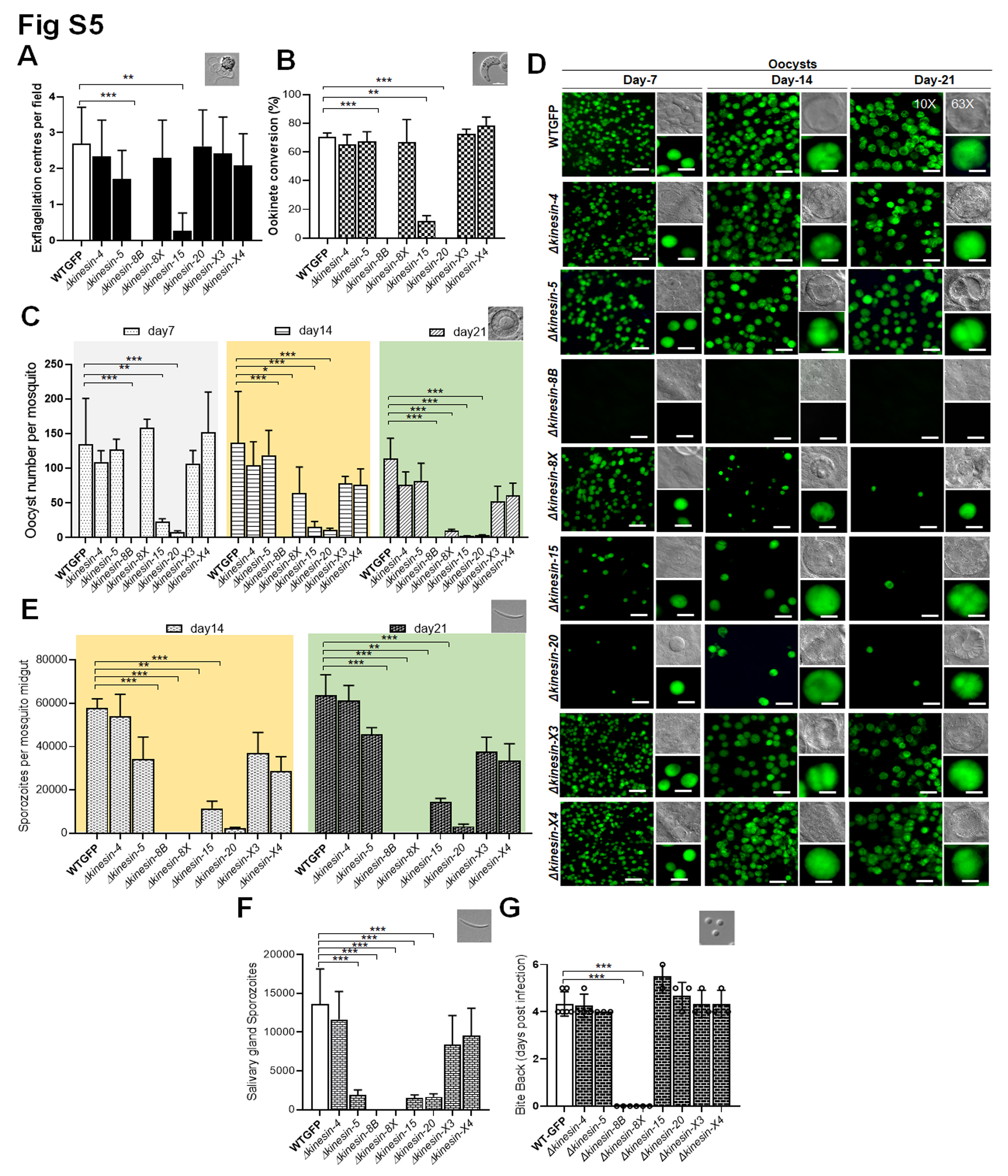

### S6

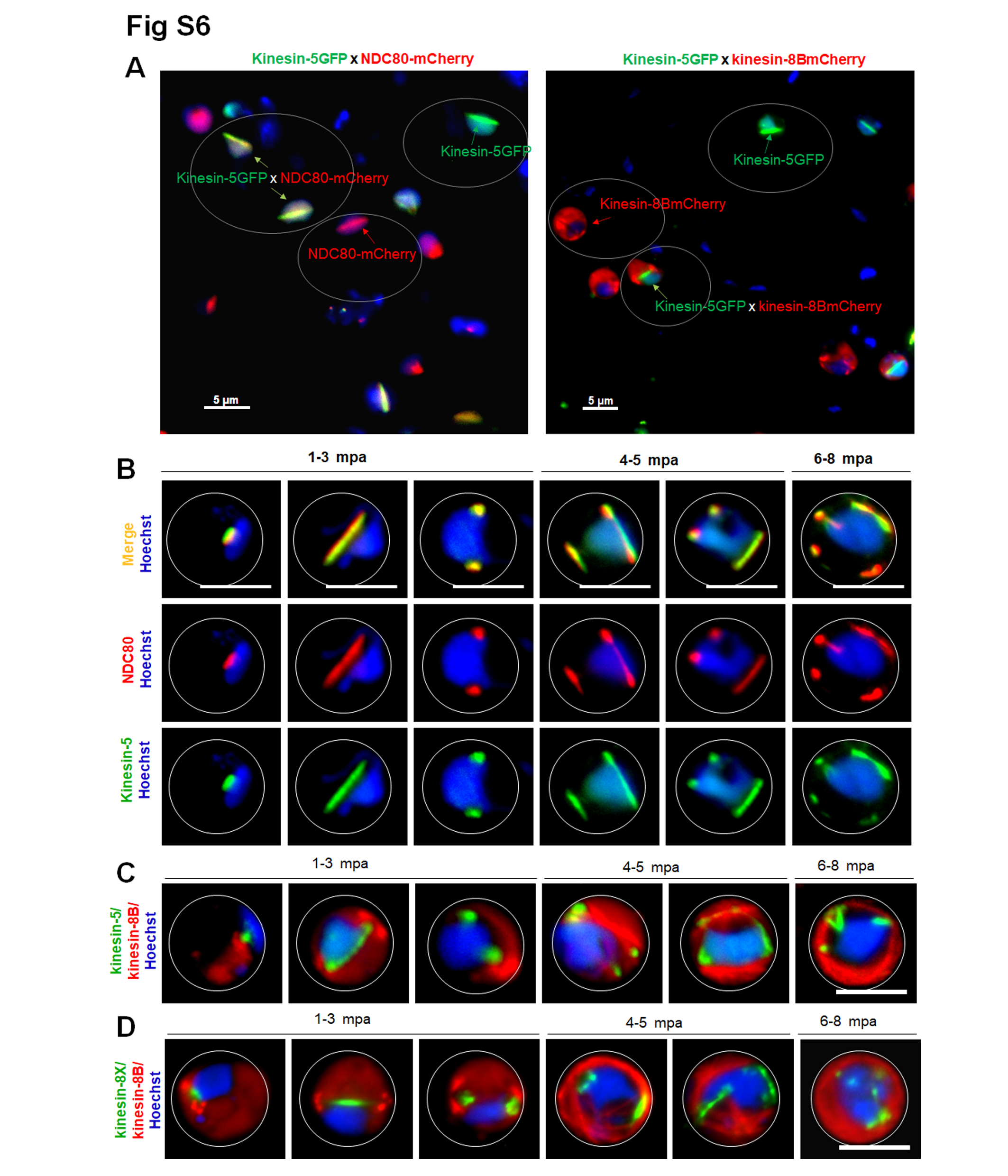

### S7

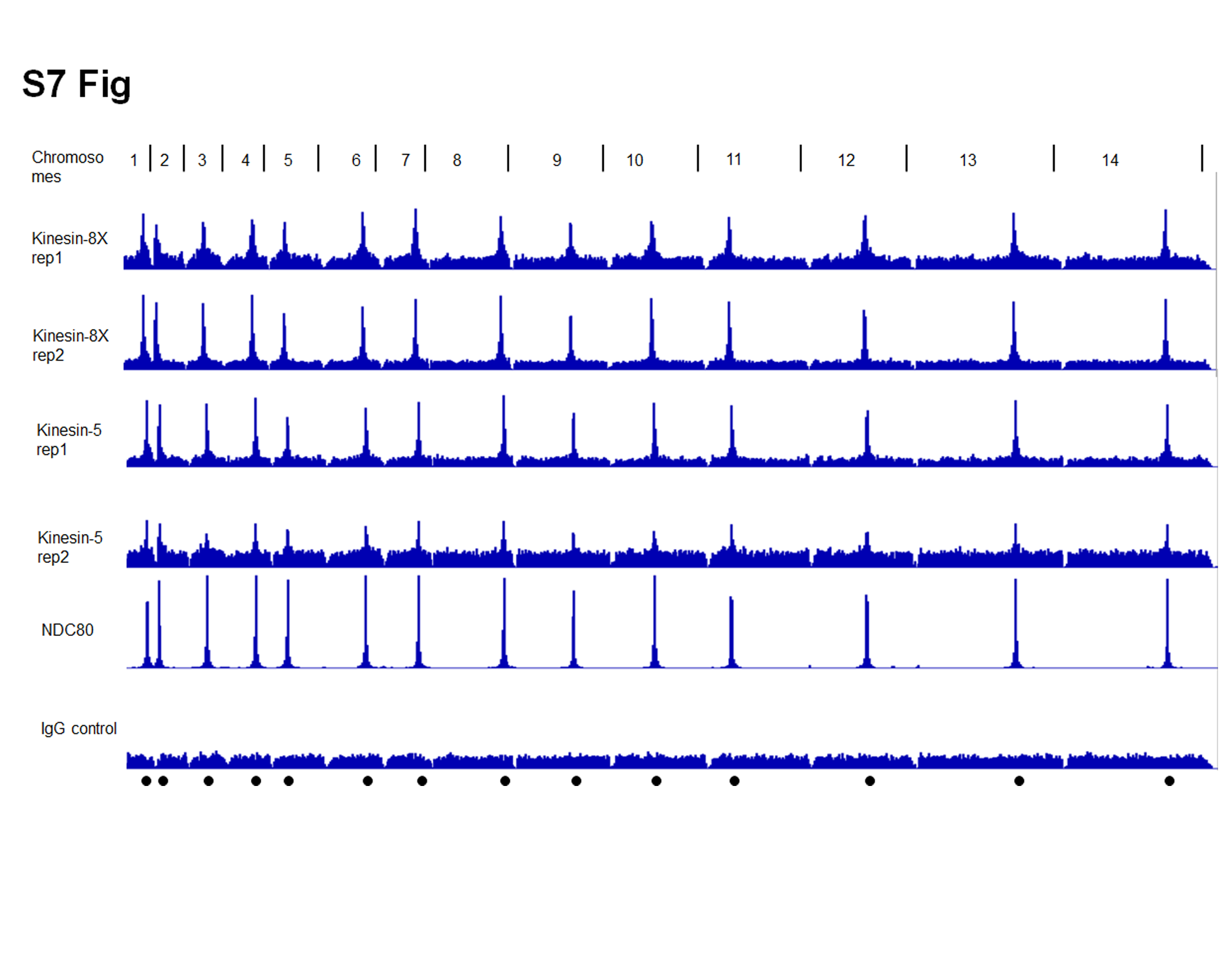

### S8

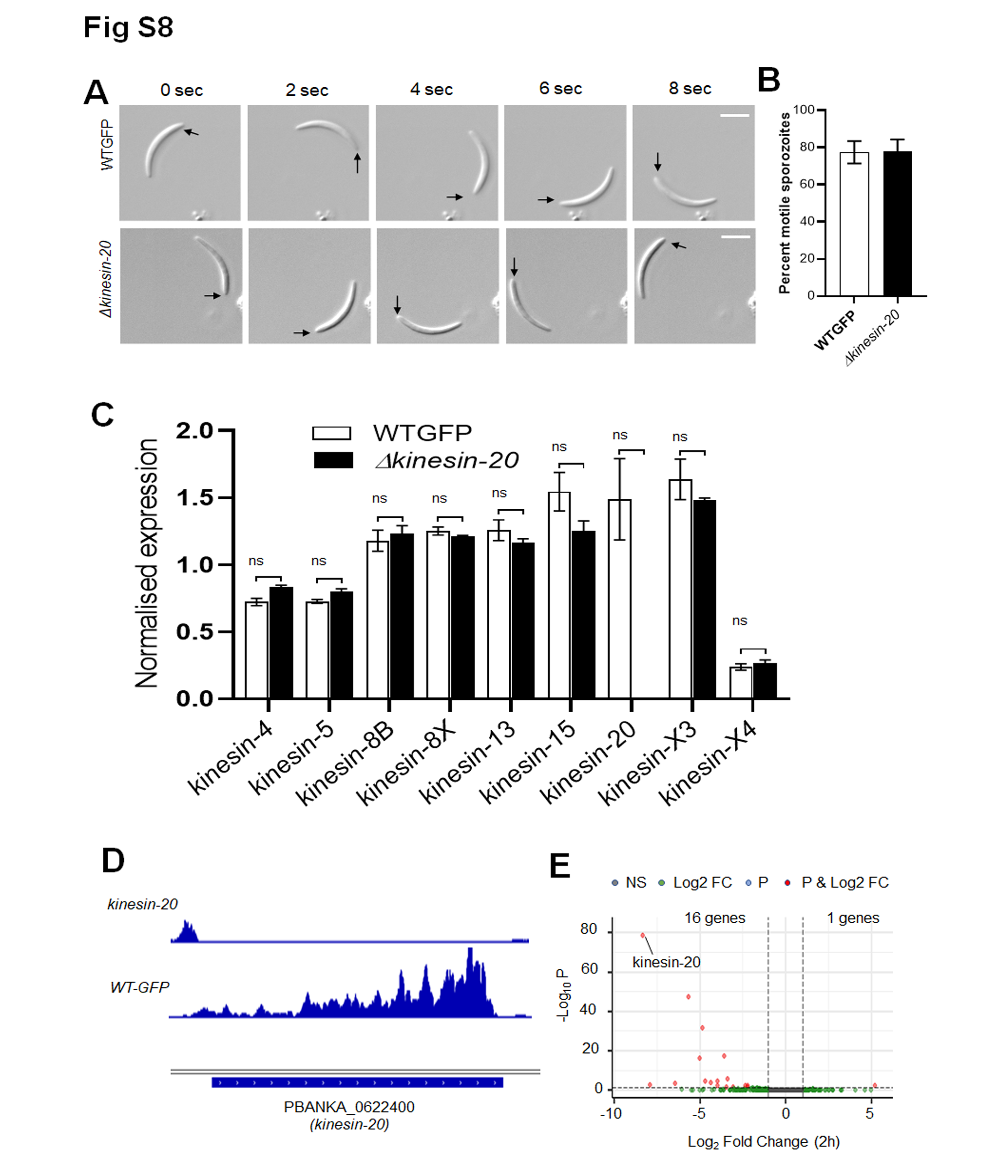

### S9

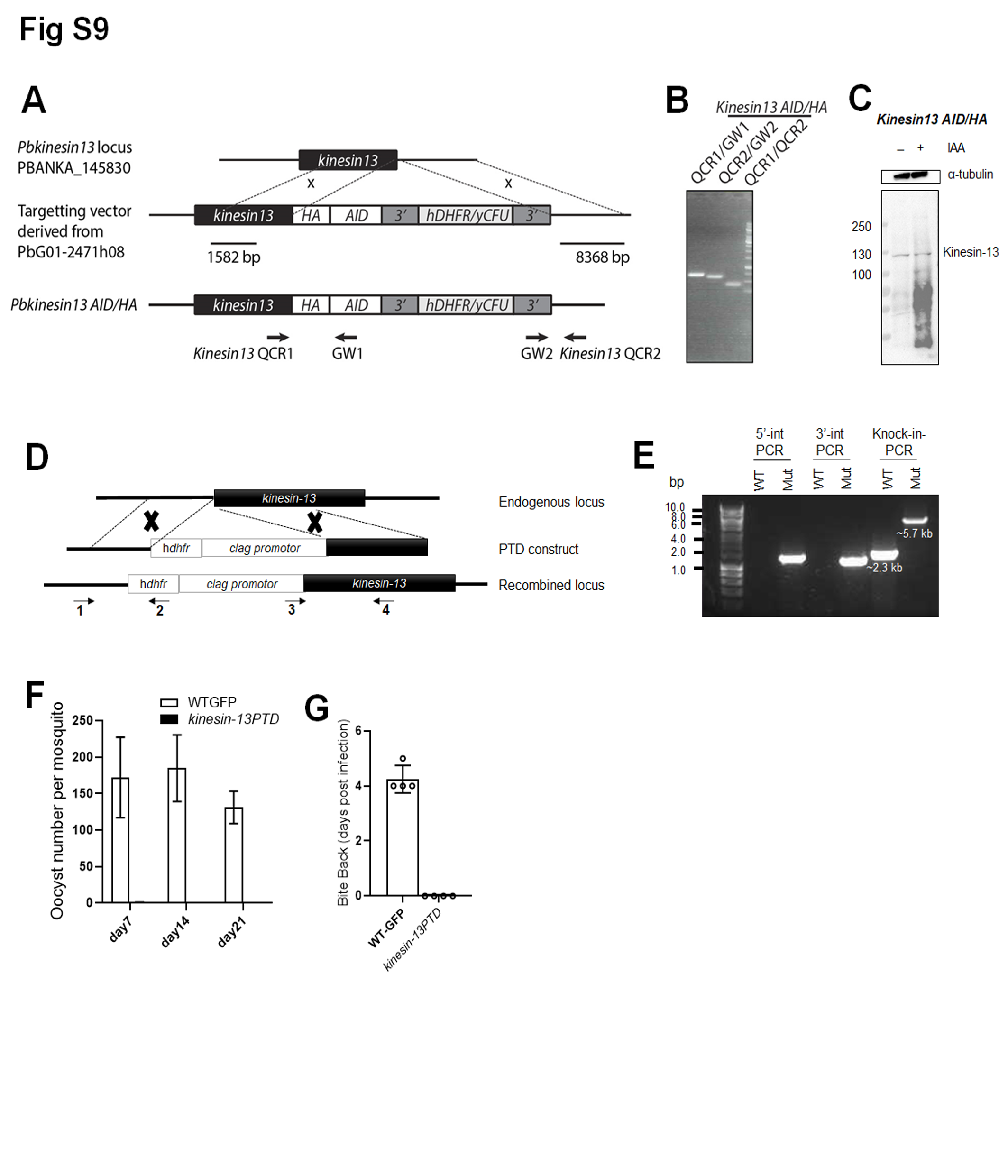
